# Long Term Study of Motivational and Cognitive Effects of Low-intensity Focused Ultrasound Neuromodulation in the Dorsal Striatum of Nonhuman Primates

**DOI:** 10.1101/2021.06.30.450605

**Authors:** F Munoz, A Meaney, A Gross, K Liu, AN Pouliopoulos, D Liu, EE Konofagou, VP Ferrera

## Abstract

Noninvasive brain stimulation using focused ultrasound (FUS) has many potential applications as a research and clinical tool, including incorporation into neural prosthetics for cognitive rehabilitation. To develop this technology, it is necessary to evaluate the safety and efficacy of FUS neuromodulation for specific brain targets and cognitive functions. It is also important to test whether repeated long-term application of FUS to deep brain targets improves or degrades behavioral and cognitive function. To this end, we investigated the effects of FUS in the dorsal striatum of nonhuman primates (NHP) performing a visual-motor decision-making task for small or large rewards. Over the course of 2 years, we performed 129 and 147 FUS applications, respectively, in two NHP. FUS (0.5 MHz @ 0.2 – 0.8 MPa) was applied to the putamen and caudate in both hemispheres to evaluate the effects on movement accuracy, motivation, decision accuracy, and response time. Sonicating the caudate or the putamen unilaterally resulted in modest but statistically significant improvements in motivation and decision accuracy, but at the cost of slower reaction times. The effects were dose (i.e., FUS pressure) and reward dependent. There was no effect on reaching accuracy, nor was there long-term behavioral impairment or neurological trauma evident on T1-weighted, T2-weighted, or susceptibility-weighted MRI scans. Sonication also resulted in significant changes in resting state functional connectivity between the caudate and multiple cortical regions. The results indicate that applying FUS to the dorsal striatum can positively impact the motivational and cognitive aspects of decision making. The capability of FUS to improve motivation and cognition in NHPs points to its therapeutic potential in treating a wide variety of human neural diseases, and warrants further development as a novel technique for non-invasive deep brain stimulation.

## Introduction

Brain stimulation techniques have long been used for the treatment of neurological disorders, and are increasingly being considered for psychiatric disorders. One of the first successful clinical uses of electrical deep brain stimulation (DBS) was to treat Parkinson’s disease (Benabid et al., 1987). Recent studies indicate that DBS in the ventral internal capsule and ventral striatum may be effective in improving symptoms of OCD (Greenberg et al., 2010; Goodman et al., 2010), while DBS in subcallosal cingulate white matter and nucleus accumbens improved some symptoms and response to antidepressants in patients with treatment-resistant major depressive disorder (Kennedy et al, 2011; Bewernick et al, 2010).

Non-invasive brain stimulation (NIBS) methods like transcranial magnetic stimulation (TMS) have also been shown to improve motor function in Parkinson’s patients (Schulz et al., 2013). TMS has been shown to help counteract the impairment of cognitive functions seen in Alzheimer’s (Boggio et al., 2011). While electrical DBS and TMS have been shown to be effective, there are some limitations. TMS is non-invasive, but is less well-localized and has limited ability to penetrate deep into the brain, reducing its applicability for subcortical structures such as the striatum (Hallet 2007). Electrical DBS can target deep structures with high spatial accuracy, but carries the risks associated with brain surgery (Mayberg et al. 2005).

Focused ultrasound (FUS) stimulation is an alternative method of non-invasive brain stimulation in which an ultrasound beam is focused onto specific regions of the brain to modulate neural activity. Studies dating back to 1929 demonstrated that FUS was able to disturb the activity of electrically active cells (Harvey 1929). FUS was first used to directly modulate brain activity when Fry and colleagues applied it to the lateral geniculate nucleus of a cat and found that it suppressed visually evoked potentials in visual cortex (Fry et al. 1959). FUS can be applied either on its own, stimulating neural activity through a mixture of thermal and mechanical effects, or with microbubbles to permeabilize the blood-brain barrier (BBB). It differs from other techniques like TMS in that it can target deep subcortical structures like the striatum. And unlike DBS, FUS is non-invasive, making it less risky and potentially suitable for a wider population (Munoz et al., 2018).

In recent years, there has been renewed interest in FUS as a clinical and scientific tool. In humans, FUS applied to the frontal-temporal cortex has been shown to slightly and temporarily reduce pain (Hameroff et al., 2013) or improve mood (Sanguinetti et al., 2020). Additionally, FUS applied to the somatosensory cortex enhances sensitivity to touch, and can even induce the sensation of touch without any external stimuli (Legon et al., 2014; Lee et al., 2016). FUS also elicits illusory visual percepts when applied to the visual cortex in human participants (Schimek et al., 2020).

While the exact neural mechanism for FUS-induced neuromodulation is still unknown, many in vivo and in vitro studies have shed light on processes that may play a role. In rat hippocampal cells, fiber volley and cell population potentials had lower amplitudes when undergoing FUS, while dendritic field potentials were enhanced (Bachtold et al., 1998). Additionally, FUS has been shown to increase sodium and calcium transients and short latency action potential firing in mouse hippocampus (Tyler et al., 2008; Tufail et al., 2010). Studies in Xenopus oocytes and C. Elegans have pointed to mechanosensitive ion channels in the brain as the target of FUS (Kubanek et al, 2018).

The current study seeks to examine the effect of FUS on motivation and decision-making when applied to the dorsal striatum of rhesus macaques. Neurophysiological studies implicate the striatum in reward-modulated sensory-motor decision making (Ding & Gold, 2013; Fan et al., 2020.) Previous studies in macaques have shown that FUS can temporarily enhance neural activity and connectivity to closely related areas in both cortical and subcortical structures (Verhagen et al., 2018; Folloni et al., 2019; Yang et al., 2021). FUS can also directly affect behavior in rhesus macaques. Several studies have showed that FUS applied to the frontal eye field alters eye movement behavior and decision-making (Deffieux et al., 2013; Wattiez et al., 2017; Pouget et al., 2020), and altered decision-making (Khalighinejad et al. 2020; Kubanek et al., 2020; Lowe et al., 2021).

A previous study from our group examined the effect of FUS on the dorsal striatum of rhesus macaques, After FUS was applied to the putamen, response speed and decision accuracy improved (Downs et al., 2017). However, in Downs et al., FUS was applied with microbubbles, which resulted in blood-brain barrier (BBB) opening, therefore the examined effects may have been a combined product of BBB opening and direct neuromodulation. Hence, the effects of applying FUS alone (without microbubbles or BBB opening) to the dorsal striatum in NHP has remained unexplored until now. The current study also addresses the long-term safety of low-intensity FUS neuromodulation. This work helps to establish parameters and protocols within which FUS can be used for beneficial effects over extended time periods without causing permanent damage.

## Materials and Methods

### Subjects

Four adult male Macaca mulatta (N, O, P, Q) weighing 9.0, 11.5, 10.5, and 8.0 kg were used in these experiments. Two (P, Q) underwent awake sonication experiments, and four (N, O, P, Q) underwent anesthetized sonication and fMRI. NHPs received vitamin enriched biscuits and enrichment toys in their cages every day. Before data collection, the two NHPs were trained for a year to perform a visual-motor decision-making task on a touch-panel display, until performance reached a consistent level. During the task, NHPs received fluid for every correct response, and continued to perform until satiated or 1000 trials had been completed. After they completed the task, NHPs were given a banana or an apple. The Institutional Animal Care and Use Committees (IACUC) of Columbia University and the New York State Psychiatric Institute (NYSPI) approved all NHP procedures.

### Behavioral Task

NHPs sat in a custom-made polycarbonate chair when performing the task (Figure 1A). The head was stabilized by means of a surgically implanted plastic post. Visual stimuli were presented on a 20-inch LCD touchscreen monitor (ELO Touch, Rochester NY) placed directly in front of the chair, so the NHPs could reach the screen. The touchscreen device had a resolution of 1,024 × 1,024 pixels, and a sampling rate of 60 Hz. Attached to the front of the chair was a vertical midline divider made of polycarbonate. This divider restricted reaching movements so that stimuli presented on the right side of the screen could only be touched by the right hand, and vice versa. The divider did not obscure vision. Both sides of the screen were equally visible at all times.

**Figure 1.**
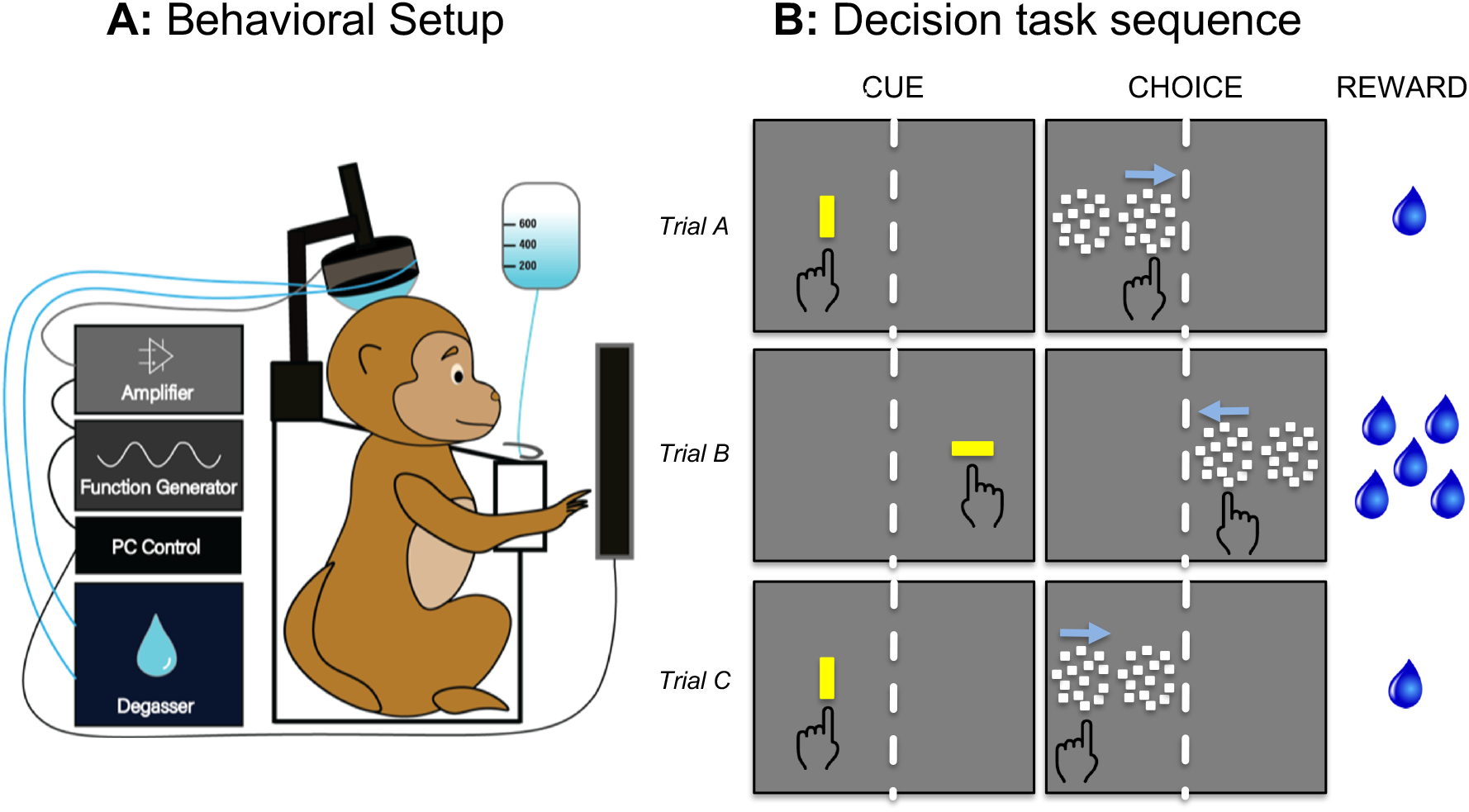
Behavioral setup, task, and session structure. **A**, Behavioral setup with NHP, chair, touchscreen, fluid delivery, transducer, amplifier, function generator, and degasser. The NHP sat in the chair so it could reach the touchscreen. The transducer was aligned to preplanned targets using a stereotaxic positioner and the Brainsight neuronavigation system. The degasser was connected to a bladder coupling system that filled the space between the transducer and the scalp. The function generator created the sonication waveform, which was amplified and input to the transducer. **B**, The task as viewed by the NHP. During the CUE period, a yellow bar prompted the NHP to initiate the trial. The orientation of the bar indicated the amount of reward given for correct performance (vertical = 1 drop; horizontal = 5 drops.) Two motion patches then appeared (CHOICE) and the correct response was to touch the patch that was moving coherently.

The task design was a two-alternative spatial forced choice (Figure 1B). When the NHP initiated a trial (Fig. 1B “CUE”), he was presented with two motion patches arranged horizontally side by side (Fig. 1B “CHOICE”). One patch contained random dots moving incoherently (zero coherence), and the other contained random dots with a motion coherence level between 0.0 and 1.0. Coherent motion could be leftward or rightward. The correct response was to touch the patch with coherent motion, regardless of the direction of motion. The coherence level was chosen randomly on each trial and all coherence levels occurred with equal probability. When both patches moved incoherently, the correct patch was designated randomly.

Each trial of the task began by presenting a vertically or horizontally oriented yellow bar (1 × 3 degrees of visual angle, 43.8 cd/m2 luminance) on the left or right side of the monitor (Fig 1B, CUE). The orientation of the bar signaled the amount of fluid reward that would be dispensed for a correct response (vertical = 1 drop, horizontal = 5 drops). By touching the oriented bar, the NHP initiated the trial. If the NHP did not touch the bar within 2.5 seconds, the trial was scored as a failure, and there was a delay of 3 seconds before the next trial began. The orientation of the bar indicated the offered reward size (1 or 5 drops) that would be delivered on correct completion of the trial.

Once the trial was initiated, the yellow bar disappeared, followed by the simultaneous presentation of 2 side-by-side patches of 100 moving dots (each dot was 0.17 deg square, luminance cd/m2.). Each patch of dots moved within a circular aperture of 10 degrees diameter, both on the same side as the screen as the original yellow bar. If the NHP touched the patch containing coherent motion, he was reinforced with drops of water (correct response). After a correct response, there was only a brief delay before the next trial began, but if the NHP touched the patch with incoherent motion there was a 3 second delay before the next trial.

Catch trials were interleaved with two-alternative choice trials. On catch trials, only the coherent motion stimulus was shown. The NHP had to touch the motion stimulus to obtain a reward. One-fifth of all trials were catch trials. The absolute coherence levels of the motion stimulus on catch trials varied from 0.0 to 1.0 in 0.1 increments, and each coherence level was presented randomly with equal probability. Coherence is signed to account for the direction of dot motion; negative coherence indicates leftward motion, positive coherence is rightward.

NHPs performed the task under 3 different experimental conditions: baseline, sham sonication, and real sonication. Under the baseline condition, NHPs completed the task without any sonication equipment set up around them. Under sham sonication, NHPs completed the task with the transducer set up on their chairs, but not activated (all of the equipment was turned on, but the FUS pressure was set to zero). Under real sonication, NHPs completed the task with the transducer set up, and a 2-minute sonication was delivered after 200 trials.

### Ultrasound Exposure

Sonication parameters were generated by a Windows 10 computer and sent to a function generator (Agilent 33220A, Agilent Technologies, Santa Clara). The output of the function generator was passed through an amplifier (E&I, Rochester, NY), which drove the ultrasound transducer (part number H-107MR, center frequency 500 kHz, focal size 2mm x 11mm; Sonic Concepts, Bothell, WA). The function generator output was monitored on an oscilloscope. The sonication carrier frequency was 500 kHz delivered in 10 msec pulses twice per second. The peak negative pressure was 200, 400, 600, or 800 kPa (kiloPascals), calibrated though a capsule hydrophone (HGL-0200, −3 dB, frequency range: 0.24-40 MHz, electrode aperture: 200mm; Onda Corp., Sunnyvale, CA). The total sonication duration was 120 sec. Sham sonications used the same setup and procedure, but the FUS pressure was set to 0.

The head was immobilized using a surgically implanted post. The transducer was positioned with a stereotaxic manipulator (Kopf, Tujunga CA). Once the transducer was positioned relative to the head, the bladder coupling system (Sonic Concepts, Bothell WA) was inflated, and coupled to the scalp with ultrasound gel. The water in the bladder was degassed using a water degassing system (WDS105+; Sonic Concepts, Bothell WA).

During sonication, the cavitation signal was recorded by a hydrophone (Y-107; Sonic Concepts, Bothell) located inside the transducer and co-axially aligned with the ultrasound focus. The hydrophone output was used to ensure adequate degassing and to monitor FUS intensity. The output was digitized by a picoscope (Pico Technology, Tyler TX).

The sonication protocol was designed to avoid auditory confounds using an offline protocol in which the sonication is delivered at a different time than the behavioral responses and fMRI are collected. Humans can detect auditory artifacts created by ultrasound, but the artifact does not appear to remain audible nor does the auditory N1 EEG persist beyond the time that ultrasound is applied (Braun et al., 2020). Hence, trials that occur before or after ultrasound application are unlikely to be affected by a peripheral auditory response. In the current study, ultrasound was applied for only 2 minutes during each behavioral session (Fig 2E). Each session was divided into three epochs: pre-sonication (200 trials), during sonication (20-30 trials), and post-sonication (up to 1000 trials). A similar approach has been used to assess “offline” persistent effects of ultrasound, which can last 1-2 hours after FUS delivery (Folloni et al, 2019; Verhagen et al., 2019, Fouragnan et al., 2019, Khalighinejad et al., 2020; Bongioanni 2021).

**Figure 2.**
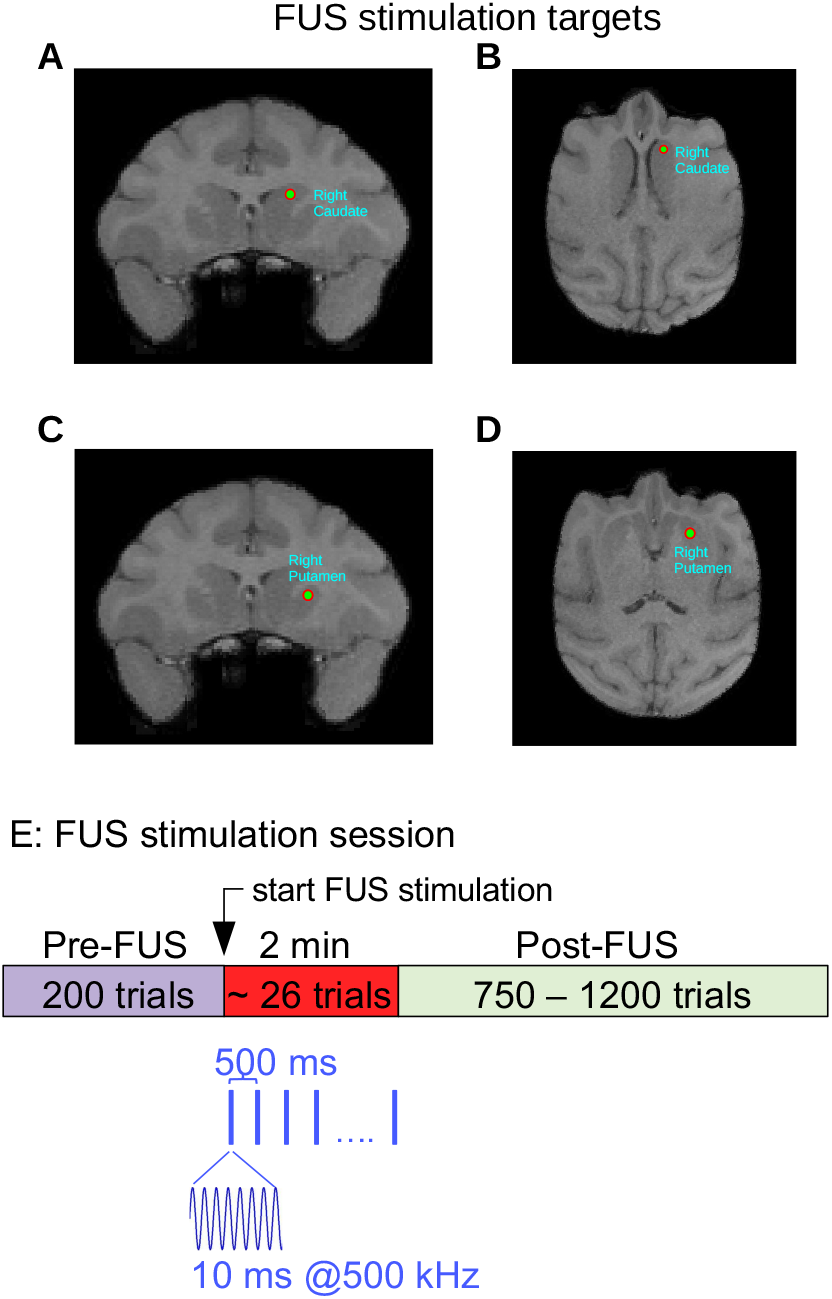
Structural MRI and Neuronavigation. **A,B**, Targeting of right caudate nucleus in NHP P. **C,D**, Targeting of right putamen in NHP P. **E**, Sonication timeline. After 200 trials, the targeted brain region was sonicated for 120 seconds. The sonication frequency was 500 kHz delivered in 10 ms bursts at the rate of 2 pulses per second. The NHP continued to perform the task during and after sonication.

### Structural MRI and Neuronavigation

A Brainsight (Rogue Research, Montreal) neuronavigation system was used to target FUS to the caudate and putamen. T1-weighted structural MRI scans (MPRAGE) were used for targeting (Figure 2A–D.) To conduct the MRIs, the NHPs were first sedated with ketamine (10mg/kg) and dexmedetomidine (0.02 mg/kg), and then anesthetized with isoflurane (1-3%) administered via endotracheal intubation. An EKG was used to monitor heart rate. The NHP was then placed in an MR-compatible stereotaxic device, with a fiducial attached to the head implant. Blankets and a warm air blower were used to maintain core body temperature. The scans were done on a 3T Siemens Prisma scanner.

Once the MRI scans were obtained, they were uploaded to the Brainsight neuronavigation system. The targets (left and right putamen, left and right caudate) and trajectories were designated for each NHP. The trajectories were selected to have a perpendicular angle with the NHP skull, in an effort to minimize ultrasound attenuation following transcranial propagation. The attenuation coefficient used was the same for both NHPs, and was derived experimentally using ex vivo NHP skulls. Prior to each experiment, the NHP was registered with Brainsight by using the pointer tool to mark each of 5 fiducials attached to the head, the same fiducial set that was used for the MRI. The orientation of the pointer tool was captured by an infrared (IR) camera, and its location was measured in reference to the subject tracker on the edge of the chair. After registration, the fiducial was removed and the transducer (Sonic Concepts, Bothell) was attached to the chair. The transducer had a tool tracker device attached to the top that allowed its orientation to be monitored by the IR camera.

At the end of the experiment, each NHP underwent a second MRI. T1 structural scans with and without gadodiamide contrast, T2 structural scans, T2-FLAIR, and susceptibility weighted imaging (SWI) scans were performed to confirm that there was no trauma.

### Functional MRI

Functional MRI scans were performed on 2 NHPs (P and O) under light anesthesia (isoflurane 0.8–1.1%, spontaneous ventilation) on a 3T Siemens Prisma scanner. A total of two post-sonication (800 kPa) and eight resting state fMRI acquisitions/runs (15 min per run) were acquired. As a control, baseline resting state functional MRI (rsfMRI) on four NHPs (O,P,Q,N) without FUS exposure and 16 acquisitions were acquired. The NHPs were scanned in the supine position while head motion was restrained with an MR-compatible stereotaxic device. In the initial phase of the anesthesia, resting state T2*-weighted gradient echo EPI functional images were acquired (TR=2000ms, TE=28.2ms, Flip angle= 70°, FOV 106×106×53mm, 64×64×32 1.65 mm^3^ isotropic voxels, 456 volumes per run) at 45 min after FUS sonication. Slices were acquired with the standard Siemens ventral to dorsal interleaved sequence. In the same session as the functional MRI, T1-weighted structural images were acquired (TR=2580ms; TE=2.81ms; FA=9°, isotropic 0.5mm resolution; FOV 128×128×60mm).

The preprocessing and analysis of fMRI data were performed using FSL (FSL 6.0.3, Jenkinson et al 2012) and Matlab (Matlab 2019b, The MathWorks, Inc., Natick, MA, United States). To correct the distortion due to B0 inhomogeneities, we also collected two short-duration (30s) echo planar images with opposite phase encoding directions. FSL-Topup was applied on both scans to generate field map images for further B0 field unwrapping (Andersson et al 2003). Brain extraction and tissue-type segmentation (white matter, gray matter and cerebral spinal fluid) were applied on structural T1 weighted images, using the BET and FAST tools of FSL, respectively (Jenkinson et al 2012). The resulting extracted brain image was linearly registered to standard D99 atlas of NHP brain (Reveley et al 2017).

The fMRI data preprocessing and first-level analysis were then performed using FSL-FEAT with procedures as follows (Jenkison et al 2012): Distortion due to motion was corrected by MCFLIRT; B0 unwrapping was applied with previously generated field map images to correct geometrical distortions; and Boundary based registration (BBR) from fMRI to structural images was performed within the GUI, using the white matter boundaries acquired from FAST of FSL. In addition, a high pass temporal filter with 100 sec cutoff was applied on fMRI data to remove the low-frequency drift; and a spatial filter with 3mm FWHM Gaussian kernel was applied on to smooth the data at each acquisition.

To calculate functional connectivity of a seed region to the rest of the brain we averaged pre-processed BOLD activity in this region and correlated it separately to the activity of all brain voxels. The seed ROI was selected as 2 × 2 × 2 mm^3^; region located at right Caudate. This procedure was repeated for all runs of the NHPs. For each voxel this rendered a set of 8 or 16 correlation values, depending on the number of functional runs with or without FUS exposure. These correlation coefficients were transformed using Fisher’s z-transformation (Fisher RA 1915) and fed into a permutation test with 5000 resamples to compare the BOLD activity of baseline rsfMRI and post-FUS rsfMRI. The resulting average seed-based correlation maps of baseline rsfMRI and Post FUS rsfMRI, as well as the difference between the two were also obtained to evaluate the FUS effects on functional connectivity.

### Audio Recording

Audio recordings were made during sonication with a lavalier microphone (Sound Professionals model SP-TFB-2-13099) placed at the entrance to the NHP’s ear canal. The microphone had a flat frequency response up to 20 kHz. The microphone output was amplified by an Etymotic Research ER-10B amplifier and digitized at 10 kHz by a National Instruments USB-6001 multfunction I/O device. A total of 36 recordings were made; six recordings ipsilateral to the transducer (2 each at 200, 600, and 800 MPa), six contralateral at the same pressures, and 24 shams (12 contralateral, 12 ipsilateral, 0 MPa). Each recording included a baseline of 2 minutes before the transducer was turned on and then 2 minutes during sonication (or the equivalent time period for shams). During recordings, all of the sonication equipment was turned on, and the NHP was alert and performing the task. During sham sessions, everything was the same as sonication sessions except that the transducer pressure was set to zero.

### Data Analysis

Matlab 8.3 with the Statistics 9.0 toolbox (Mathworks, Natick MA) was used for data analysis, with all statistical equations in the Mathworks format. Response times (RT) were analyzed with multivariate ANOVA to determine significance and a generalized linear model regression (using the Matlab glmfit function) to measure effect size. The explanatory variables were 1. an identifier for each NHP subject (P or Q), 2. motion coherence (−1.0 to 1.0), 3. Amount of offered reward (1 or 5 drops), 4. presence of sonication. FUS pressure and sonicated hemisphere (ipsilateral or contralateral to responding hand) were included in some analyses

Performance accuracy was analyzed with multivariate ANOVA and logistic regression. The same explanatory variables used in the response time GLM were also used in this analysis. Psychometric functions (accuracy vs coherence) were estimated by using Gaussian process regression (GPR; Rasmussen & Williams 2006). Gaussian processes (GPs) is a powerful non-parametric probabilistic framework for performing regression and classification. GPR is a highly flexible non-linear estimation technique that can be used to inference for overdispersed and/or missing data. GPR is performed by estimating the extent to which every observation covaries with every other, given some prior metric for comparing the distance between observations along the dimensions of interest. Each observation influences the estimate for other ‘nearby’ observations (e.g. that occur at similar times or have similar coherence) more than observations that are distant (Lucas et al. 2015). Although a full Gaussian process model can be computationally prohibitive to fit, we take advantage of the “expectation propagation” approximation (Tolvanen et al. 2014) implemented in the GPstuff toolbox (Vanhatalo et al. 2013) to accelerate estimation.

## Results

The goal of the current study was to determine if low intensity FUS affects behavioral performance when directed to the dorsal striatum (caudate and putemen) in NHPs performing a motivated decision-making task. We hypothesized that FUS directed to the anterior caudate and putamen might be associated with changes in reaching accuracy, motivation, decision accuracy, and reaction time.

Both NHP were trained on the task shown in Figure 1B for one year before any real or sham sonications were performed. During this year of training, they attained stable, asymptotic performance levels. The data reported in the current study were collected after the initial year of training. NHP P completed a total of 169 behavioral sessions (129 with real sonication, 40 sham), with an average of 1400 trials per session. NHP Q completed a total of 188 behavioral sessions (147 with sonication, 41 sham), with an average of 1402 trials per session.

Sonications were spread out over a period of 2 years. The large number of sonications did not impair either NHPs’ ability to perform the task. On the contrary, performance actually improved slightly. Over the course of the entire experiment, the average performance for NHP P improved from 78.0% correct responses during the first 20 sonication sessions to 79.3% during the last 20 sonications (t-test p=0.2), and for NHP Q from 84.4% to 85.7% (t-test p=0.02). These small improvements are evidence against the hypothesis that long-term application of FUS led to a degradation in cognitive performance.

### Effects of FUS on Touch Precision

Touch precision is a measure of motor performance. Here, touch precision is defined as the dispersion of the first point of contact the NHP made with the screen after the motion stimulus appeared. Figure 3A shows the effect of sonication on touch precision for reaches to the motion stimulus in three example sessions (one each with sham, putamen, or caudate sonication) for NHP P. This analysis was restricted to trials in which a single motion patch was presented, to avoid confounding reach accuracy with decision uncertainty. Only post-sonication trials in each session (trial numbers 230 to 1030) were used for analysis.

**Figure 3.**
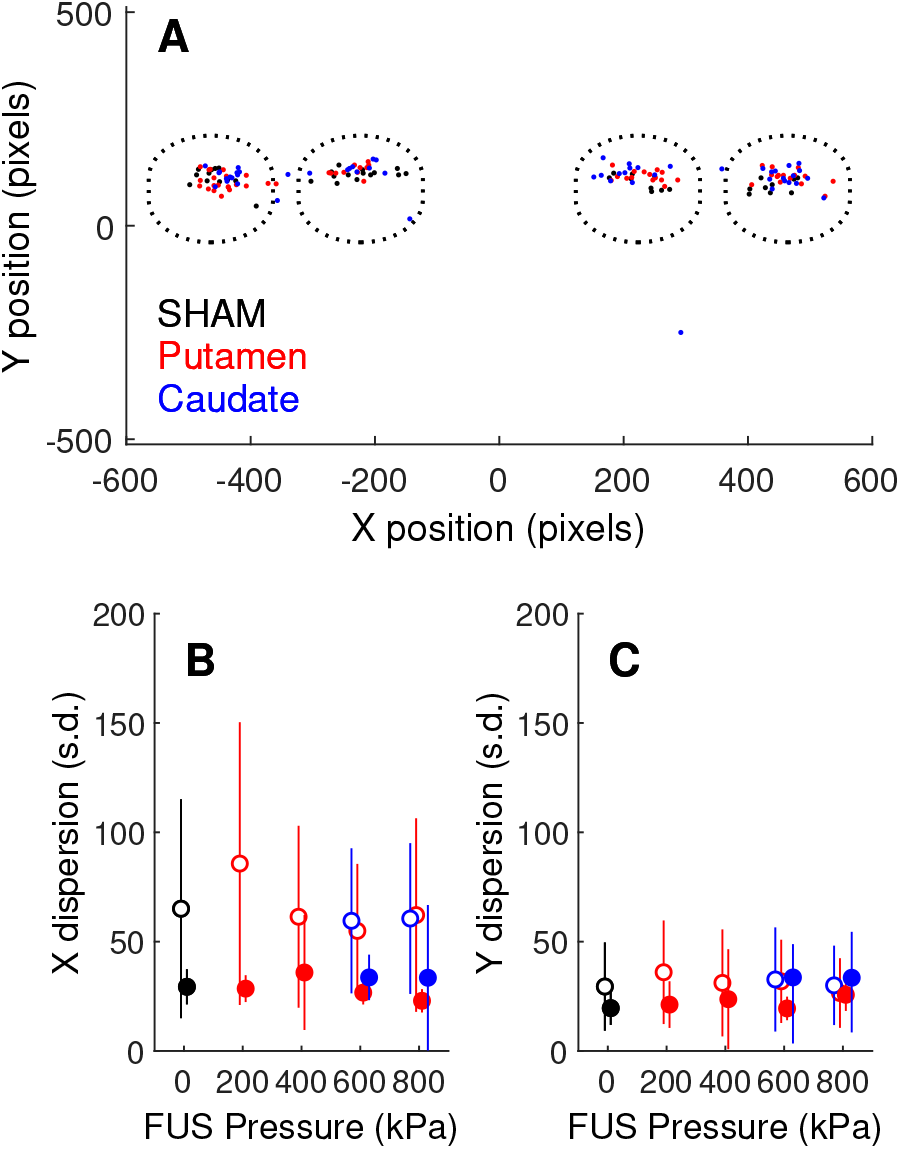
Effects of FUS on touch precision. **A**, X and Y initial touch coordinates for three sessions (one each of sham, putamen sonication and caudate sonication) for NHP P. The dotted circles indicate the approximate size and locations of the motion stimuli. **B**, X-axis dispersion (standard deviation of initial touch coordinates) as a function of FUS pressure (0 = sham). Open symbols are NHP P, closed are NHP Q. Data point are slightly shifted horizontally for clarity. **C**, Y-axis dispersion of touch coordinates (same conventions as **B**).

Touches were clustered near the center of the motion target. Clustering was similar in trials with or without sonication, indicating that touch precision was not impacted by FUS to the caudate or putamen. It was expected that sonication of the putamen might have a greater motor effect than sonication of the caudate, as the putamen is the striatal component of the motor “loop” (Parent & Hazrati 1995), but sonication of neither brain area had an effect on the NHPs’ ability to make spatially accurate reaching movements.

The standard deviation of the initial X and Y touch coordinates relative to the center of the motion stimulus was taken as a measure of initial touch variability. The average X and Y dispersion is plotted as a function of FUS pressure in Fig. 3B,C. A one-way ANOVA was used to test the significance of the effect of FUS pressure on touch dispersion across sessions. The results are shown in Table 1 along with a measure of effect size (eta-squared = percentage of total variance accounted for by variation in FUS pressure from 0 to 800 kPa). In no case did FUS pressure account for more than 10% of the variance in initial touch dispersion.

**Table 1.** ANOVA for effect of FUS pressure on touch precision.

| NHP | Region | EV | X dispersion |  |  | Y dispersion |  |  |
| --- | --- | --- | --- | --- | --- | --- | --- | --- |
|  |  |  | p-value | df | effect size | p-value | df | effect size |
| P | putamen | pressure | 0.27 | 4, 118 | 4.3% | 0.62 | 4, 118 | 2.2% |
| P | putamen | pressure | 0.87 | 2, 73 | 0.4% | 0.85 | 2, 73 | 0.5% |
| Q | caudate | pressure | <0.01 | 4, 136 | 10.0% | 0.20 | 4, 136 | 4.3% |
| Q | caudate | pressure | 0.64 | 2, 73 | 1.2% | 0.05 | 2, 73 | 8.1% |

### Effects of FUS on Motivation

The primate dorsal striatum is strongly associated with reward-motivated visual-motor behavior (Ding & Hikosaka, 2006). Both the likelihood and speed with which goal-oriented actions are initiated have been considered to be behavioral indicators of motivation (Tourè-Tillery & Fishbach, 2014). The motion detection task was designed so that each trial was self-initiated by the NHP. This provided an opportunity to assess the effect of FUS on motivation by quantifying the NHP’s willingness to initiate trials. Trials were initiated by touching the cue (a yellow bar) that was presented at the start of each trial (Figure 1B). The orientation of the bar indicated the amount of reward that the NHP would receive for a correct response on that trial (vertical = 1 drop, horizontal = 5 drops). If the NHP did not touch the cue within 2.5 seconds, the trial ended without a reward and was scored a “failure.” The rate at which failures occurred was roughly 4 times lower when the cue signalled a larger reward size (Fig. 4A vs. 4B.) Within each session, the failure rate increased with the number of trials completed (Fig. 4A,B), which is correlated with satiety. The dependence on reward size and satiety suggest that failure rate is a valid index of motivation.

**Figure 4.**
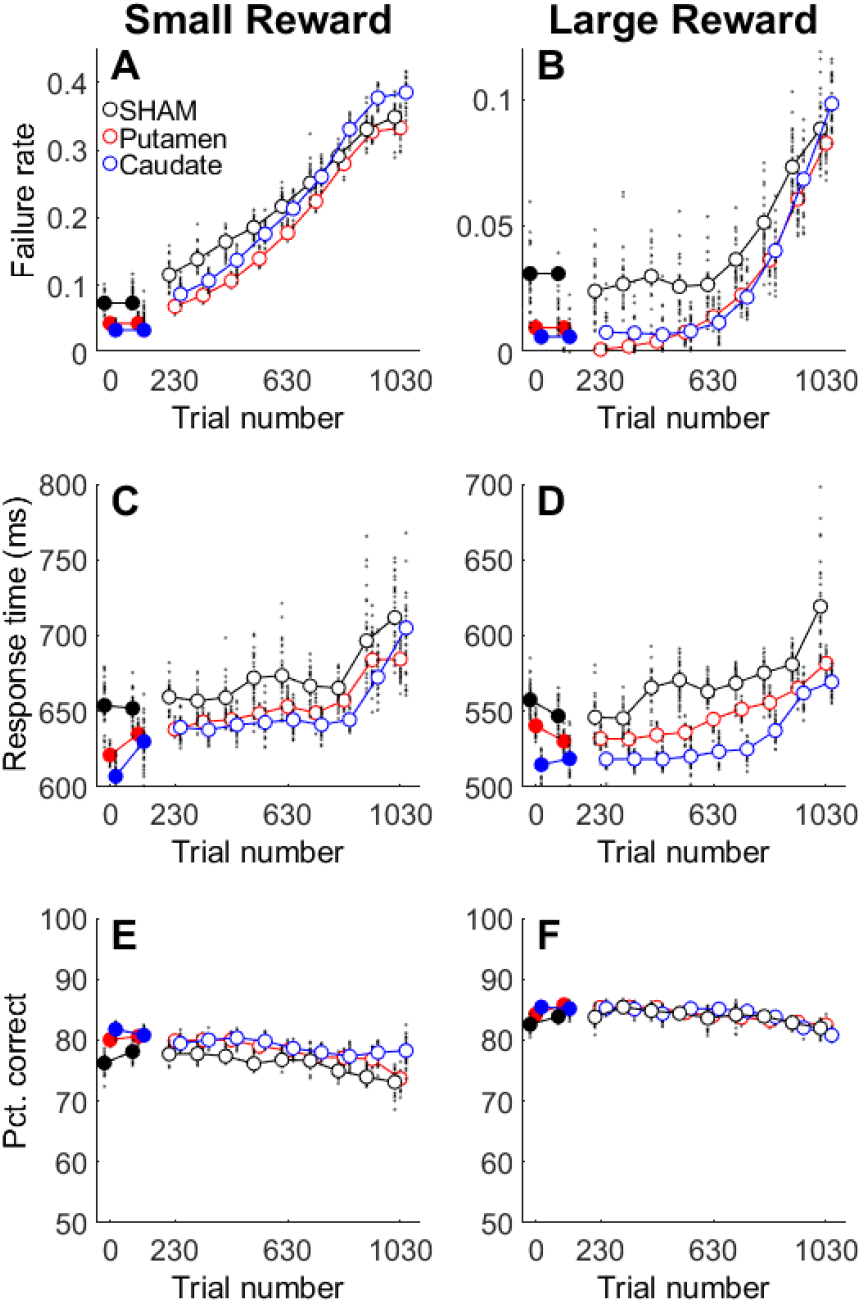
Effects of FUS on motivation. Pre-FUS trials (filled symbols) are grouped into two ranges (0-100, and 100-200 trials before FUS). Post-FUS trials (open symbols) are grouped into 100-trial bins from 230-1030 after FUS. Data points are slightly offset horizontally for clarity. The small black points are bootstrap estimates of the mean. **A** Effect of sonication on failure rate for low reward trials. **B**, Effect of sonication on failure rate for high reward trials. Note that the y-axis scale is different for A and B. **C**, Effect of sonication on response time for low reward trials. **D**, Effect of sonication during high reward trials. Note that the y-axis range is different for C and D, but the scale is the same. **E**, Effect of sonication on overall decision accuracy for low reward trials. **F**, Effect of sonication on decision accuracy for high reward trials.

The failure rate for trials following FUS administration was quantified by counting the number of failures in successive 100-trial intervals in each session, spanning pre-FUS (trials 1-200) and post-FUS (230-1030) trials. These counts were converted to the probability of failing to initiate any given trial. These probabilities were then averaged across sessions. Results are shown in Figure 4 for low (Fig. 4A) and high (Fig. 4B) reward trials. There were differences in the pre-FUS failure rates, but also differences in the post-FUS timecourse that depended on sonication treatment.

Failure rate during post-FUS trials depended significantly on reward size, and whether FUS was applied to the striatum. Table 2 shows the results of 2-way ANOVAs with the following explanatory variables: treatment (FUS or sham) and reward size (1 or 5 drops). The dependent variable was the normalized failure rate, i.e. the number of failures to initiate a trial after the FUS was applied (trial number 230 to 1030) minus the failure rate on pre-FUS (trials 1-200). N was calculated as the (number of sessions) × (treatment levels) × (reward levels). Effect size is reported as partial eta-squared (effect sum-of-squares divided by effect + error sum-of-squares), which is an estimate of the variance accounted for by each explanatory variable. In the caudate, the effect of FUS was comparable in magnitude to the effect of reward size, whereas, in the putamen, the effect of FUS was much smaller than reward (Table 2).

**Table 2.** ANOVA for effect of reward size and FUS treatment on failure rate.

| NHP | Region | EV | ANOVA F | ANOVA P | N | Effect size |
| --- | --- | --- | --- | --- | --- | --- |
| P | putamen | FUS | 28.6 | <0.0001 | 472 | 0.06 |
| P | putamen | reward size | 95.3 | <0.0001 | 472 | 0.17 |
| P | caudate | FUS | 39.5 | <0.0001 | 168 | 0.29 |
| P | caudate | reward size | 55.9 | <0.0001 | 168 | 0.32 |
| Q | putamen | FUS | 7.1 | <0.01 | 556 | 0.03 |
| Q | putamen | reward size | 89.3 | <0.0001 | 556 | 0.19 |
| Q | caudate | FUS | 57.3 | <0.0001 | 164 | 0.41 |
| Q | caudate | reward size | 47.1 | <0.0001 | 164 | 0.40 |

Response times may also be an indicator of motivation. For trials that were initiated by the NHP, the response time (RT) was defined as the time elapsed between the initial appearance of the cue stimulus and the first touch registered by the touchscreen. RT was faster for larger rewards, but tended to become slower over the course of the behavioral session as the NHP became satiated with rewards (Fig. 4C,D). RT thus appears to be inversely correlated with motivation. Table 3 quantifies differences in RT that were correlated with sonicated region (caudate or putamen), presence of FUS treatment, and reward size for both NHP (P and Q). The GLM coefficients (beta) are estimates of effect size scaled in milliseconds. Large rewards were associated with reduction of response time of approximately 100-150 msec. FUS Sonication also modestly reduced response time relative to sham controls, with caudate sonications resulting in slightly greater reductions than putamen.

**Table 3.** ANOVA and GLM for effect of sonicated region, FUS, and reward size on initial response time.

| NHP | EV | ANOVA F | ANOVA P | N (trials) | GLM beta (msec) |
| --- | --- | --- | --- | --- | --- |
| P | region | 6.9 | 0.001 | 86053 | -1.2 |
| P | FUS | 61.9 | <0.0001 | 86053 | -14.7 |
| P | reward size | 11542.7 | <0.0001 | 86053 | -155.4 |
| Q | region | 607.0 | <0.01 | 93142 | -22.4 |
| Q | FUS | 123.4 | <0.0001 | 93142 | -18.6 |
| Q | reward size | 10397.3 | <0.0001 | 93142 | -98.6 |

Overall decision accuracy (percent correct responses on choice trials) is another potential indicator of motivation. Accuracy tended to be highest at the beginning of each session and then fell off as the NHP became satiated (Figure 4E,F). Decision accuracy was roughly 5% higher and showed less variability for large reward trials than for small rewards (compare Fig. 4E and 4F.) As was the case for failure rate and initial response time, the dependence on reward size and satiety strongly suggests that overall decision accuracy is a valid measure of motivation.

There were only modest differences in overall accuracy between sonication conditions (sham vs. caudate vs. putamen). Table 4 provides the results of an ANOVA. Effect size is measured as partial eta-squared (proportion of total variance accounted for by each explanatory variable. N is number of sessions × reward levels.) The effect of reward size on overall accuracy was roughly an order of magnitude stronger than the effect of FUS sonication.

**Table 4.**
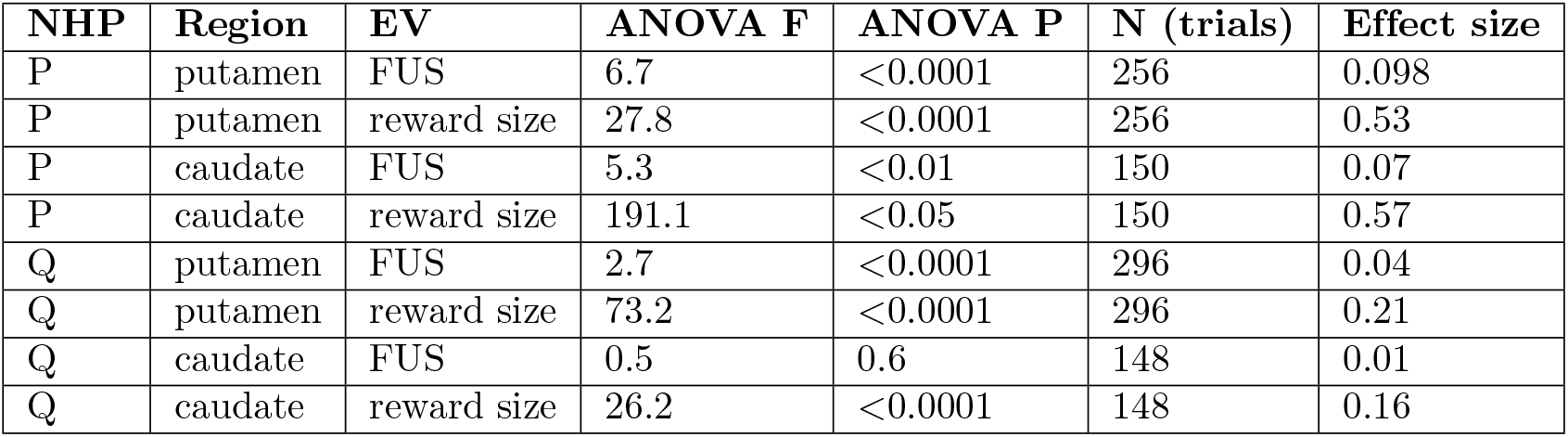
ANOVA for effect of FUS and reward size on overall decision accuracy.

### Effects of FUS on performance: accuracy and response time as a function of motion strength

Performance on the motion detection task was quantified in terms of accuracy (percent correct) and response time (time from motion stimulus onset to first registered touch response) as a function of motion coherence for choice trials. Coherence takes into account the direction of dot motion; negative coherence values indicate leftward motion, positive coherence corresponds to rightward motion. Post-sonication behavior (trials after trial number 230 in each session) during FUS sessions was compared to the same trial range in randomly interleaved sham sessions. Performance (Figure 5 A,B) and response time (Figure 5 C,D) were affected mainly by motion coherence and reward size. NHP responded faster and more accurately as absolute coherence increased, and were also more accurate but slower when a larger reward was offered. FUS sonication had significant but small effects on accuracy and response time. These effects did not depend strongly on the sonicated region (putamen or caudate) or hemisphere. The relative effect size for each explanatory variable (reward size, FUS pressure, sonicated hemisphere) was estimated using the odds ratio for decision accuracy, and by a generalized linear model for response time (Figure 5 E,F). Session number was included as an explanatory variable to account for long-term trends in accuracy and response time over the course of the entire experiment. The explanatory variables were scaled to the same range (0 to 1) so that the magnitude of the regression beta values could be directly compared. For example, the effect of FUS pressure on accuracy, while statistically significant, was much smaller than the effect of reward.

**Figure 5.**
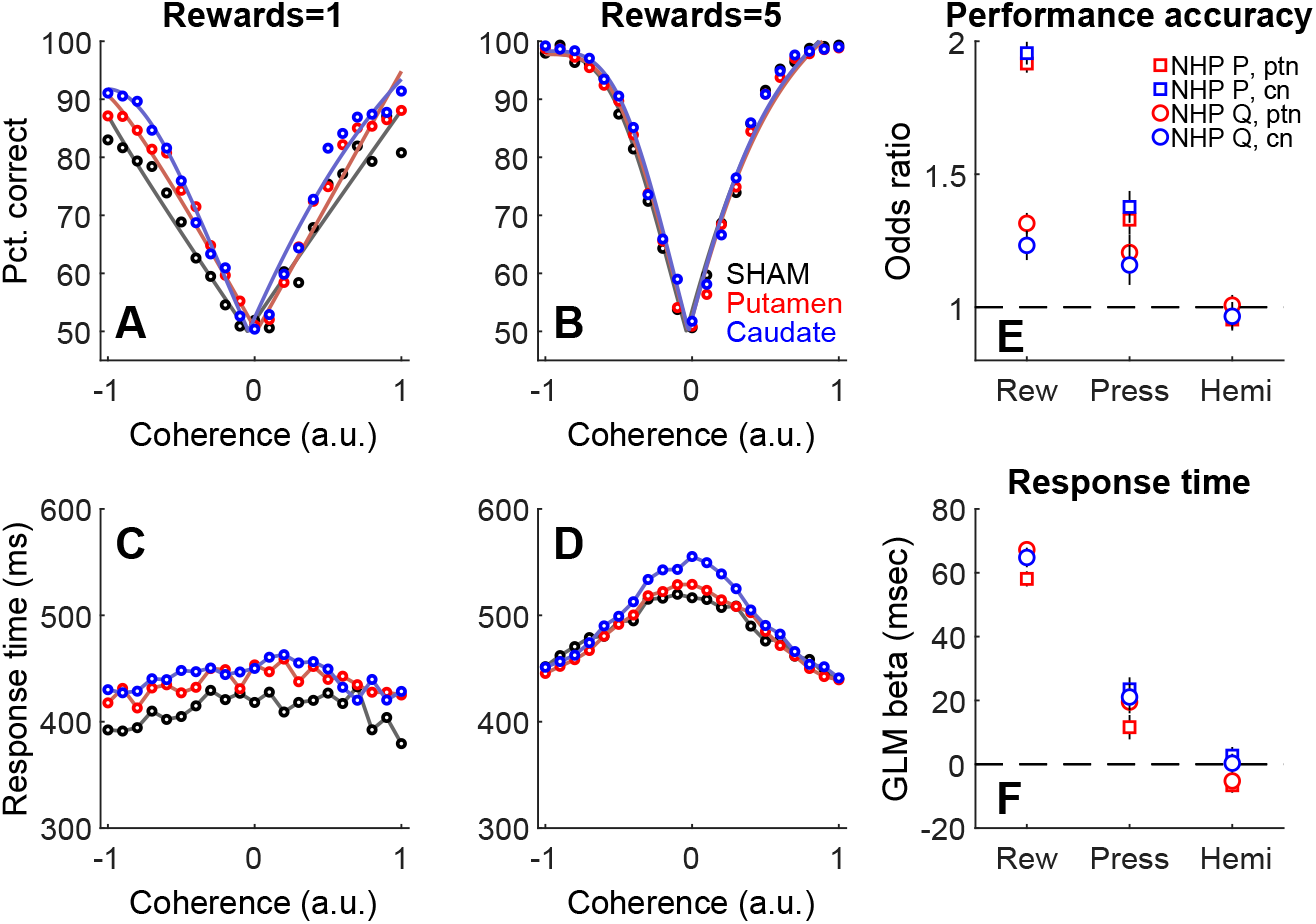
Effects of FUS on decision-making performance. Effects of sonication on decision accuracy (**A,B**), and response time (**C,D**) for NHP P. Reward size is indicated in the titles above each column. Colors indicate condition: sham (black), putamen sonication (red), or caudate sonication (blue). Open circles are overall averages across sessions. Solid curves in top row are fits of psychometric functions to mean accuracy data. E, Effect size (odds ratio) of reward (Rew), sonication pressure (Press) and sonicated hemisphere (Hemi) on performance accuracy for both NHP and both sonication targets (ptn = putamen, cn = caudate nucleus). Error bars are ±1 standard deviation. F, Effect sizes (GLM beta) of reward, sonication pressure, and sonicated hemisphere on response time for both NHP and targets. Same legend as E.

**Figure 6.**
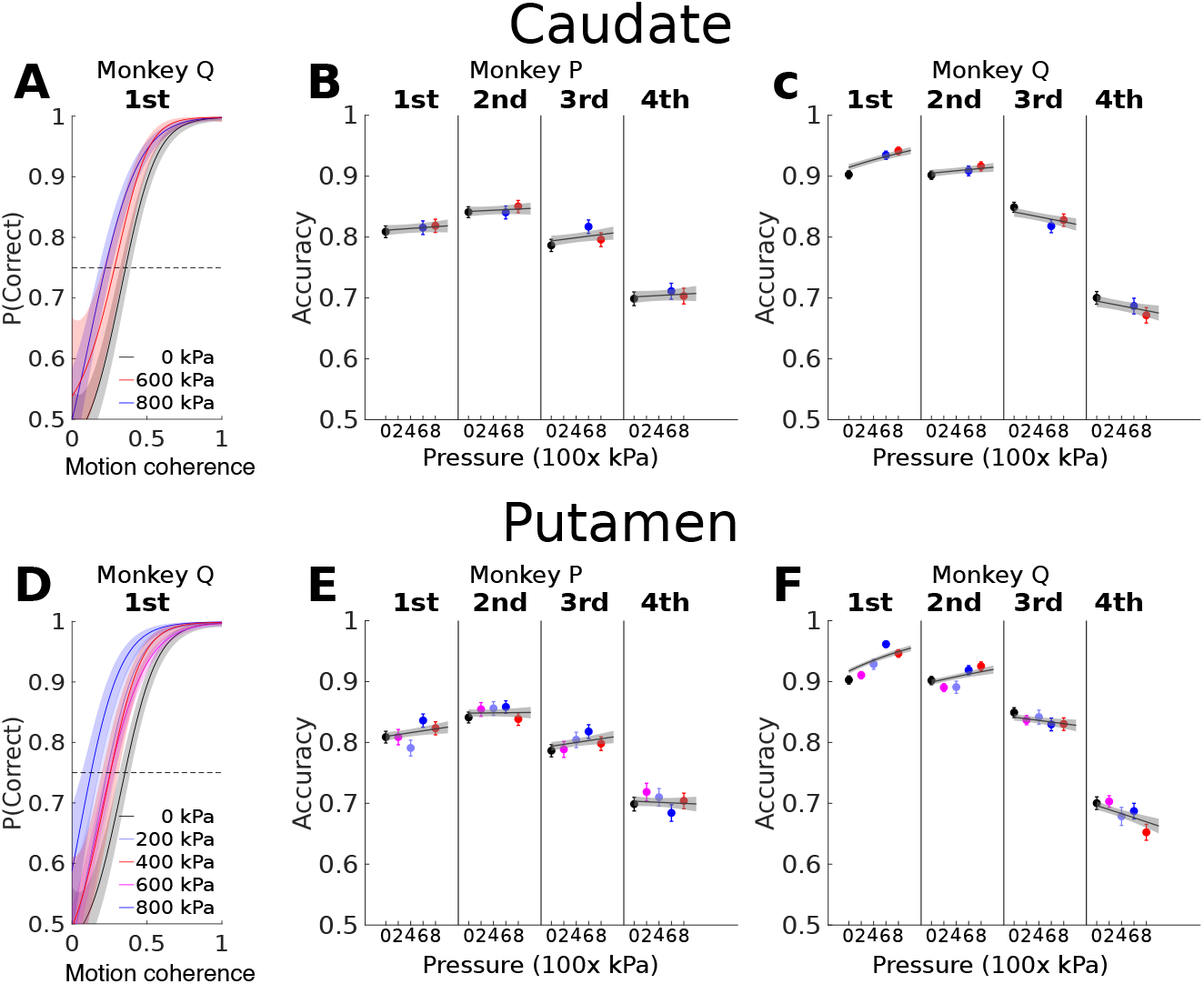
Performance (overall percent correct) divided into response time quartiles, as a function of sonicated region and pressure. **A,D**, Psychometric functions using non-parametric estimation (gaussian process regression) applied to the performance accuracy in the first quartiles in the distribution of response time in the case of FUS stimulation in caudate and putamen at pressure ranging from 0 (sham) to 800 kPa in monkey Q. **B,C**, the accuracy estimation with 90% confidence intervals in accuracy estimation for each response time quartile for monkeys P and Q when FUS stimulation was applied to caudate, the line correspond to the linear relation with the pressure by applied GLM, the gray region is the 95% confidence interval of the regression estimation. **E,F**, the same procedure as in B,C, but applied to putamen stimulation.

Response time variability can provide insight into decision processes. A widely used approach for analyzing RT variability is to divide RT distributions into quantiles (Ratcliff & McKoon, 2008). Choice trials were sorted into four RT quartiles and decision accuracy was calculated in each quartile for each NHP, region, and FUS pressure. For both NHP and sonicated regions, accuracy was highest for faster responses (1st and 2nd RT quartiles), and tended to increase with FUS pressure (Figure 5) For slower responses (3rd and 4th quartiles), accuracy was lower overall and the dependence on FUS pressure was not consistent across NHP or regions.

The effect of FUS pressure on performance accuracy in different response time quartiles was fit with a generalized linear model. The GLM beta weights segregated by NHP, sonicated region (caudate or putamen), and response time quartile are shown in Table 5.

**Table 5.**
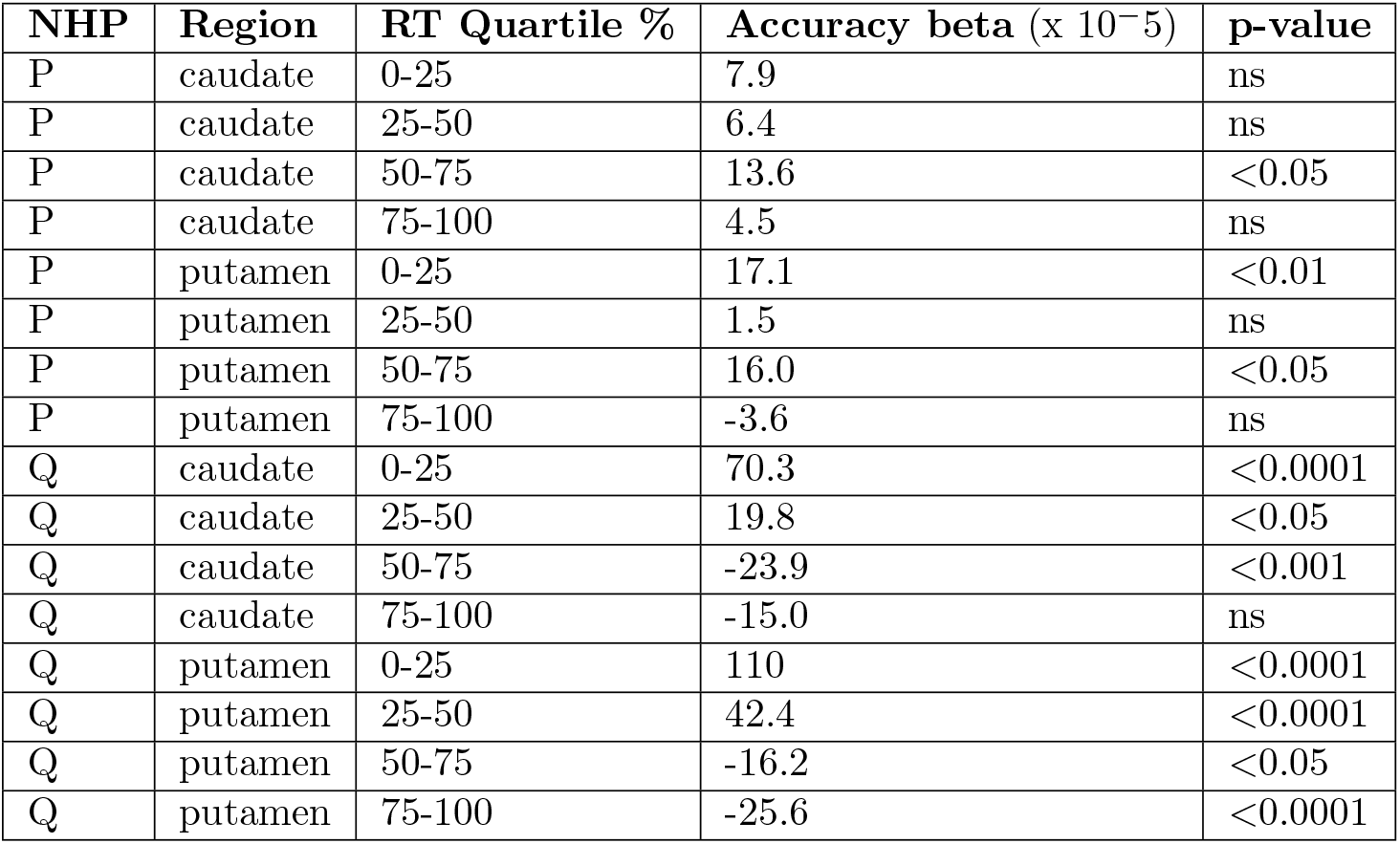
GLM for effect of FUS pressure on performance accuracy in response time quartile.

### Audio Recordings and Performance during Sonication

Although the behavioral data reported above were collected “offline,” i.e. before and after sonication, there may still be concerns about effects mediated by auditory stimulation. To address this, we made audio recordings during putamen sonications with a microphone placed at the entrance to the NHP’s ear canal. Figure 7A shows the amplitude spectrum (up to the Nyquist frequency of the digitizer) of the microphone output during real and sham sonications. The thin lines are the mean amplitude, averaged over 6-12 sessions. Shaded regions are the standard deviation of the mean. The inset plot shows the spectrum for low frequencies (0-10 Hz). For reference, the pulse repetition frequency of the sonication was 2 Hz. Across the entire spectrum, the recorded sound pressure during real sonications did not differ from that during shams.

**Figure 7.**
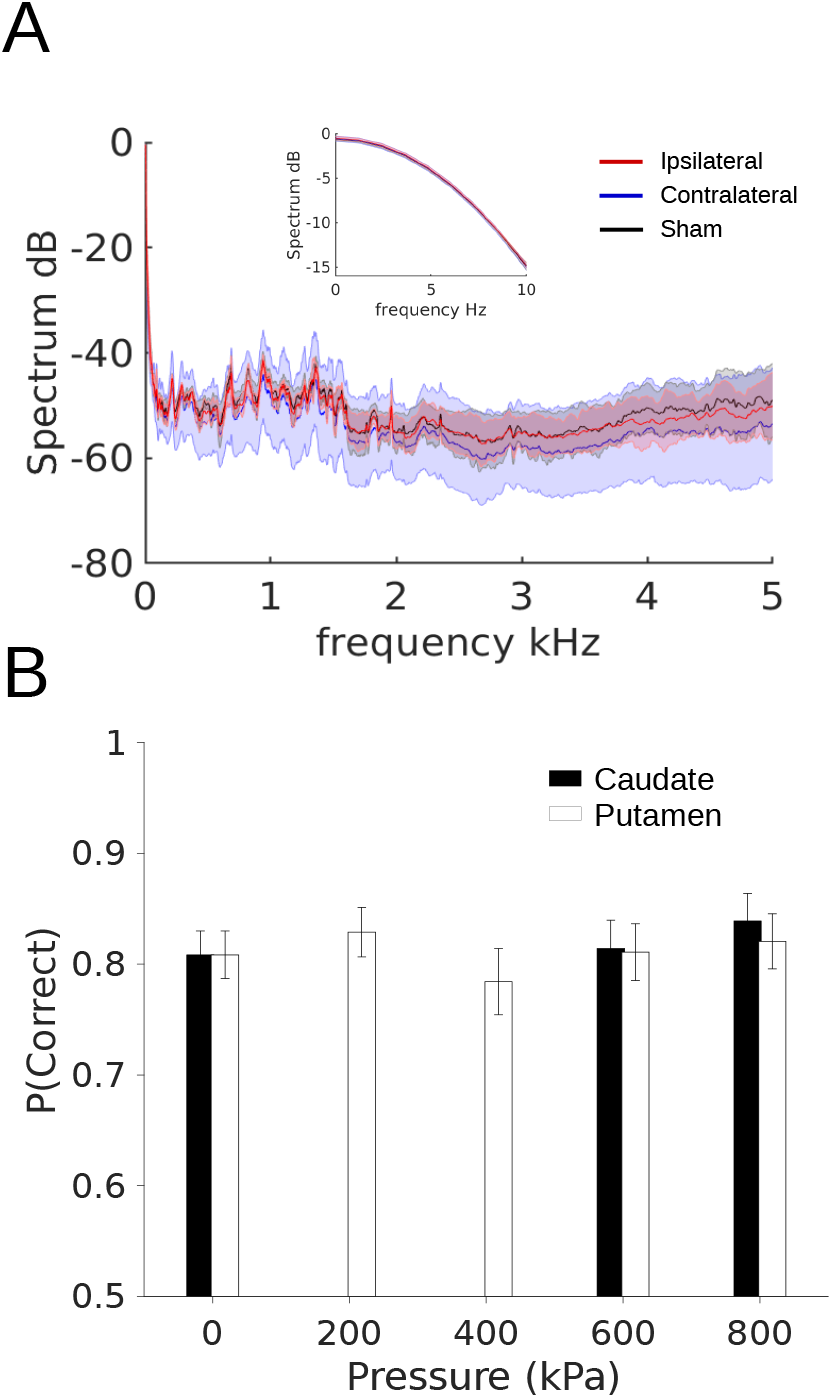
Audio recordings and performance during sonication. **A**, Frequency spectrum of recordings from microphone place ipsilateral (red) or contralateral (blue) to transducer, or during sham sonication (black). **B**, Overall percent correct for trials during sonication. Error bars are standard deviation of a beta distribution of the proportion of correct and incorrect responses.

To test if behavior was disrupted by FUS sonication, behavioral performance (overall percent correct) during sonication is shown in Figure 7B. Due to the smaller number of trials (30 per session), the data were modeled with a beta distribution. The bars and whiskers are the mean and variance of the beta distributions in each condition. Performance during real sonications was slightly better than shams. However, none of the pairwise comparisons showed a significant difference between real and sham sonications (chi-squared < 3.2, p > 0.05, df = 1).

### Structural MRI

FUS has the ability to heat and displace neural tissue, which could lead to permanent lesions. FUS might also disrupt the blood supply or weaken blood vessels, leading to infarct or hemorrhage. Structural MRI’s were performed under anesthesia before the first sonication and after the last (Figure 8). T1-weighted, T2-weighted, and susceptibility weighted (SWI) images were obtained. For post-FUS T1 (Fig. 8, middle column), the images were combined with T2-weighted images to enhance contrast. T2 and SWI images were not obtained pre-FUS. SWI is sensitive to blood, which shows up as regions of hyperintensity.

**Figure 8.**
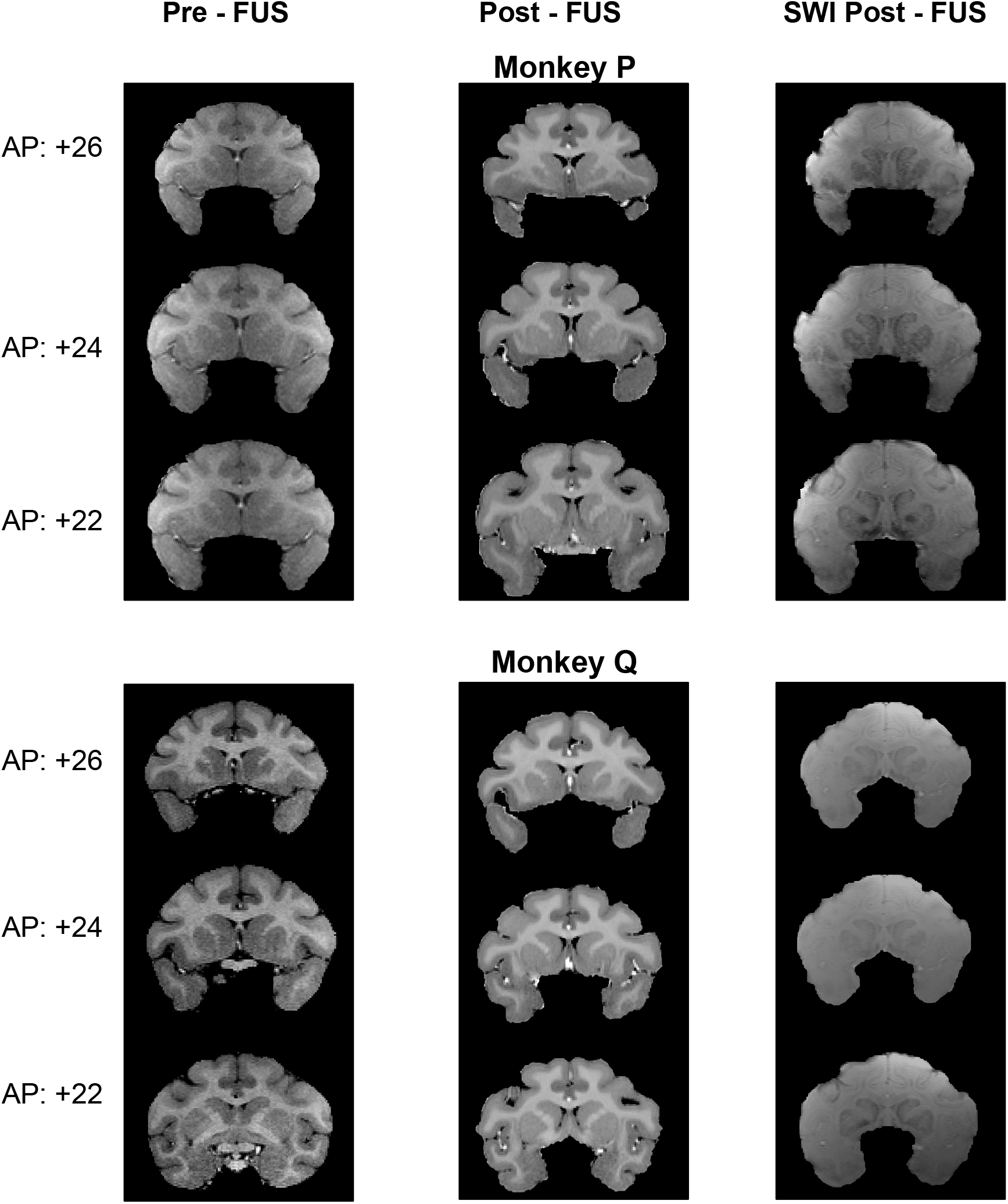
Structural MRIs at the start and end of the experiment for NHP P and Q. **Left**, “Pre-FUS” T1-weighted MRI before the NHP had experienced any sonications. **Middle** “Post-FUS” T1-weighted MRI after all sonications had been completed. **Right** “Post-FUS” susceptibility-weighted MRI.

Neither NHP P or Q had been treated with FUS prior to this study. Hence, the pre-FUS T1 images show their normal brain anatomy. No evidence of chronic or acute trauma was visible in the post-FUS images, nor were there significant changes compared to pre-FUS MRI. These results do not rule out the possibility that trauma arose and was resolved prior to the end the study. However, if this happened, it did not leave any lasting evidence. Both NHP are currently being used in follow-up studies.

### Functional Connectivity

Seed based correlation maps of rsfMRI and post FUS sonication rsfMRI, as well as the difference map are shown in Fig.9 A-C overlaid on the standard template (D99 Atlas, Reveley et al, 2018) of NHPs in sagittal, horizontal and coronal planes respectively.

**Figure 9.**
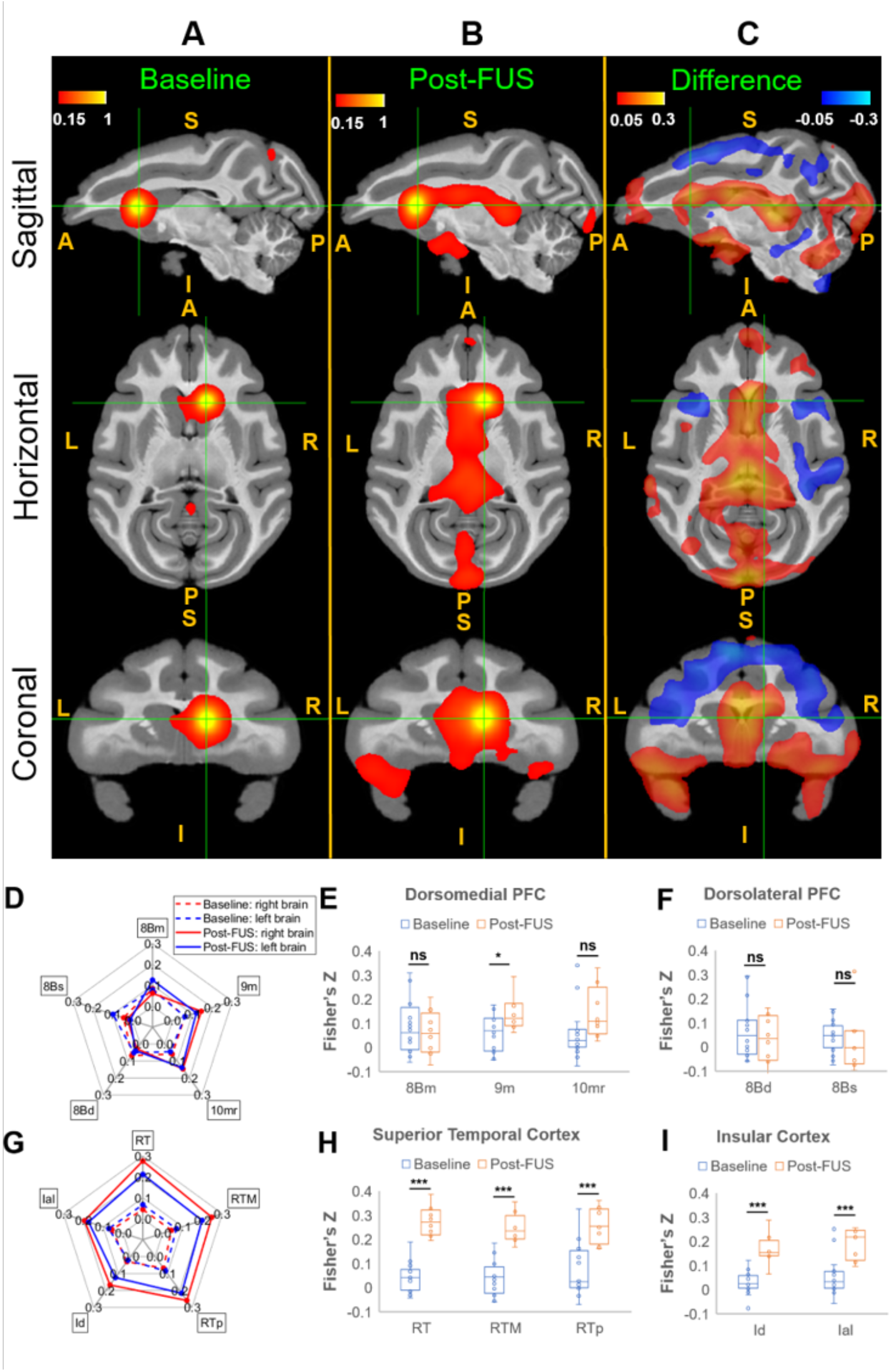
Average seed-based correlation maps. **A**, Baseline rsfMRI (16 runs). **B**, Post-FUS rsfMRI (8 runs), **C**, The difference (B-A) in sagittal, horizontal and coronal planes (S: Superior; I: Inferior; A: Anterior; P: Posterior; L: Left; R: Right). The seed ROI was chosen as the FUS sonication target-right caudate, shown in the green cross in A,B, and C. All the correlation data were transformed to Fisher’s Z-scores. Mean Correlations between the selected regions from Dorsomedial Prefrontal Cortex (dmPFC), Dorsolateral Prefrontal Cortex (dlPFC), Superior temporal cortex (STC) and Insular Cortex (IC) and the seed ROI at baseline (dotted) and post-FUS (solid) were shown in the spider-plots (**D**: dlPFC and dmPFC and **G**: STC and IC), and statistical analysis in the right hemisphere (**E**: dmPFC; **F**: dlPFC; **H**: STC; **I**: IC) between Non-FUS and FUS were performed using the permutation test with 5000 resamples, with ns denoting p>0.05, * denoting p<0.05 and ***denoting p<0.001. Abbreviations: 8Bm - Area 8B medial, 9M - Area 9 medial, 10mr - Area 10 medio-rostral, 8Bd - Area 8 dorsal, 8Bs - Area 8B arcuate sulcus, RT - rostrotemporal cortex, RTM - medial rostrotemporal, RPp - rostrotemporal polar, Id - dysgranular insula, Ial - lateral agranular insula.

With FUS exposure in the right caudate, an increased mean correlation between the right caudate seed ROI and the dorsomedial prefrontal, superior temporal, and insular cortex was observed. A decreased correlation between right caudate and the dorsal premotor cortex, dorsolateral prefrontal cortex, and ventrolateral prefrontal cortex were found. We selected these specific brain regions and performed permutation tests between the baseline resting state fMRI (16 acquisitions/runs) and post-FUS resting state fMRI (8 acquisitions/runs).

The mean correlation coefficients are expressed in terms of Fisher’s Z at selected regions in prefrontal cortex, and temporal and insular cortex, and are shown in Fig. 9D, and Fig. 9G respectively. The corresponding results of permutation tests with 5000 permutations are shown in Fig. 9 E-F and H-I, respectively.

The increased correlations between right caudate and the selected regions in the superior temporal cortex and insular cortex due to FUS exposure showed statistically significant differences between baseline and post-FUS results (p<0.001). However, the results do not show strong statistical significance in the PFC, with p-values greater than 0.01. The selected brain regions in the PFC showed no significant correlation (p>0.05 for regions 8Bm,10mr 8Bs and 8Bd) or weak significance (0.01<p<0.05 for 9m).

## Discussion

The potential of low-intensity focused ultrasound for neuromodulation, first recognized in the 1950’s (Fry 1959), has recently garnered strong renewed interest. Along with this increasing enthusiasm, a number of concerns have arisen, not least among these being the long-term safety of repeated applications of ultrasound in the brain. To address this, we performed an extended study in which two rhesus macaques received FUS exposure in the dorsal striatum once or twice a week over the course of two years, resulting in 129 and 147 sonications, respectively. These numbers represent 5-10 times more sonications than previously published studies. To compare with the current literature, Table 6 provides sonication parameters, number of subjects, brain targets, and average number of sonications per subject in all of the studies of which we are aware that were performed with awake, behaving NHP. In the current study, we assessed the effects of FUS exposure behaviorally and with structural MRI. There was no detectable motor or cognitive impairment, nor damage visible on MRI, consistent with past studies (Bystrysky et al, 2011).

**Table 6.**
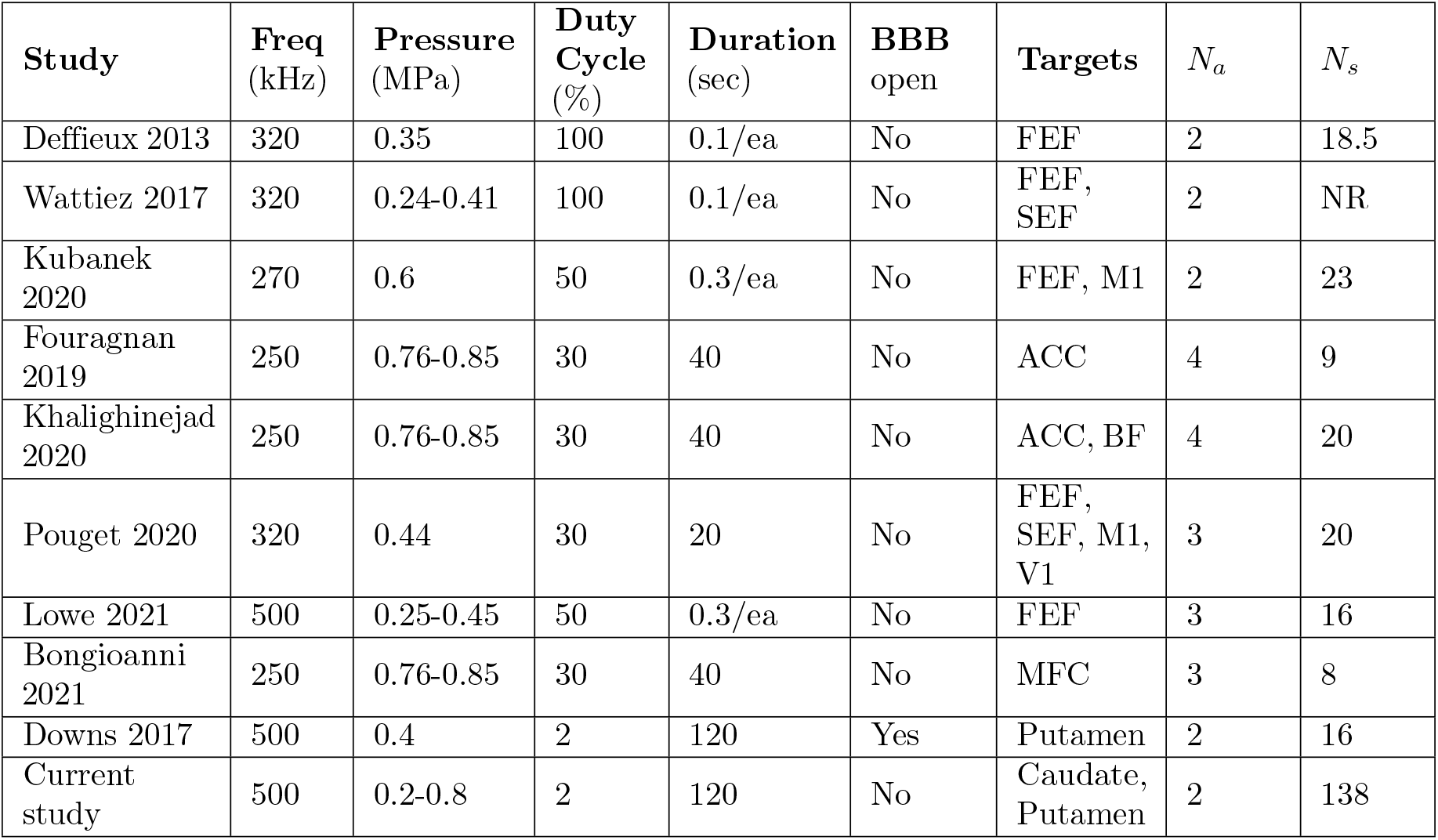
FUS neuromodulation studies in awake, behaving NHP. Abbreviations: FEF – Frontal Eye Filed, SEF – Supplemental Eye Field, ACC – Anterior Cingulate, BF – Basal Forerain, M1 – motor cortex, MFC – Mediat Frontal Cortex, NR – not reported, *N_a_* - number of animals, *N_s_* - avg. number of sonications.

The choices of brain regions and cognitive task were motivated by prior work that established a role for the dorsal striatum in reward-motivated behavior and decision-making (Hikosaka et al., 2014; Ding & Gold, 2013). The goal of the current study was not to test hypotheses about the function of the dorsal striatum, nor to compare sonication of dorsal striatum to sonication of other structures, but to determine if focused ultrasound in the dorsal striatum would impair or facilitate performance of a task suited to that structure. This would help determine the potential of focused ultrasound for therapeutic applications. FUS exposure resulted in increased motivation and small improvements in decision accuracy, compared to sham controls. The effects on motivation, measured by the speed and probability of initiating a trial, were modest but statistically significant. The effects on decision-making performance were much milder. However, it is clear that long-term sonication did not impair decision-making.

Previous work has demonstrated that FUS applied to the frontal eye field (FEF, Deffieux et al., 2013) can impact saccadic movements and also modulate activity in a connected cortical area (Wattiez et al., 2017). Kubanek et al. (2020) found that FUS can bias decisions when applied to the FEF but not to primary motor cortex. Other studies found temporary changes in cortical activity and connectivity after FUS application in rhesus macaques (Verhagen et al., 2018; Folloni et al., 2019). However, only a few studies have shown effects on motivation or cognition (Bongioanni et al., 2021.)

The current study is the first to show that FUS alone (without microbubbles) applied to the caudate and putamen of rhesus macaques affects motivational and cognitive aspects of behavioral performance in a motivated decision-making task. The NHP were more willing and faster to initiate the task during FUS sessions as compared to sham. They also performed the task with slightly higher accuracy. Response times were a bit slower after FUS exposure. However, the improvement in accuracy was most consistent for trials with fast response times. These improvements were FUS dose dependent, raising the possibility of larger effects with higher pressures, higher duty cycles or lower center frequencies than were tested here. However, the safety of applying higher pressures and duty cycles to the brain must also be considered.

A previous study by our group (Downs et al. 2017) demonstrated improved performance on a motion discrimination task several hours after the administration of FUS with microbubbles that resulted in blood-brain barrier (BBB) opening. In Downs et al., (2017), NHP showed improved accuracy along with shortened response times, a pattern that points to improved decision efficiency. In the current study, NHP were more accurate but slower to respond, a pattern that is more consistent with a speed-accuracy trade-off. Hence, FUS neuromodulation of the dorsal striatum without BBB opening may have qualitatively different effects that FUS-mediated BBB opening. This is in contrast to the results of a recent study in senescent mice (Blackmore et al., 2021) in which FUS was directed to the dentate gyrus of the hippocampus with and without BBB opening. In that study, BBB opening after FUS with microbubbles led to recovery of long-term potentiation (LTP). FUS alone, with no BBB opening also restored LTP, but was even more effective than FUS-mediated BBB opening.

While these results and past studies point to FUS altering neural activity and behavior, one main limitation of FUS research is that the exact mechanism of action is unknown. There is evidence that FUS does not act through the same mechanism as electrical stimulation. A recent study determined that while the cortex became refractory to further stimulation for several seconds after FUS, the same effect was not seen after electrical stimulation, suggesting two distinct modes of action on the brain (Gulick et al., 2017). One past study found that FUS application to slices of rat hippocampus (dentate gyrus) reduced the amplitude of fiber volley and cell population potentials, while increasing dendritic field potentials (Rinaldi et al., 1991; Bachtold et al., 1998). Additionally, a 2008 study by Tyler et al. found FUS application triggers calcium and sodium transients in slices of mouse hippocampus. Recently, it was discovered that FUS modulated the activity of specific mechanosensitive potassium and sodium channels in Zenopus oocyte cells, but it is yet to be determined if these results translate to neurons (Kubanek et al., 2016). Additionally, it is unclear how long the changes brought on by FUS last, with some studies reporting effects lasting milliseconds and others reporting up to 40 minutes (Hameroff et al., 2013; Deffieux et al., 2015).

In addition to altering neuronal function at the cellular level, FUS may act at the circuit-level. This may be revealed by changes in functional connectivity even when FUS is applied outside the MRI scanner (Verhagen et al, 2019). Here, we found that FUS directed to the caudate nucleus significantly increased correlated fMRI-BOLD activity in multiple cortical areas, including dorsomedial prefrontal cortex, anterior insula, and superior temporal cortex. Resting state connectivity between the striatum and dorsomedial prefrontal cortex has been described anesthetized macaques (Yacoub et al., 2020). Anterior insula has been implicated in decision-making under uncertainty in humans (Grinband et al., 2006) and NHP (Wittmann et al., 2020), as has superior temporal cortex (Paulus et al. 2001.) While these changes in functional connectivity are plausibly related to activation of known decision-making networks, it should be noted that the current results do not establish a causal link between changes in functional connectivity and behavior.

FUS has the advantage of being non-invasive and able to reach deeper brain structures. For these reasons, FUS shows promise as a potential treatment for neurological and psychiatric disorders. A few studies have been published showing focused ultrasound to be effective in treating Parkinson’s and Alzheimer’s, but in these cases ultrasound was either used with microbubbles or as a neurosurgical technique (Lipsman et al., 2006; Martinez-Fernandez et al., 2018; Magara et al., 2014). Beisteiner et al (2019) did not use microbubbles but still measured a memory improvement in AD patients. Overall, low-intensity FUS can be a relatively low-risk and effective method of deep brain stimulation.

## Acknowledgments

We thank the following lab members who helped with data collection: Saani Borge, Grant Spencer, David Freshwater, and Krystian Loescher. Ray Lee supervised fMRI data acquisition. Jack Grinband provided advice on fMRI data analysis. Greg Jensen provided comments on a draft of the manuscript. This work was generously supported by NIH grant R01-MH112142 to VPF and EK, R01-EB009041 and R01-AG038961 to EK, and a NARSAD Young Investigator Award to FM.

## Notes

### Competing Interest Statement

The authors have declared no competing interest.

